# Plexin-Bs enhance their GAP activity with a novel activation switch loop generating a cooperative enzyme

**DOI:** 10.1101/796748

**Authors:** Zhen-lu Li, Jeannine Müller-Greven, SoonJeung Kim, Luca Tamagnone, Matthias Buck

**Author notes:** equal contributions (Zhen-lu Li, Jeannine Müller-Greven).

## Abstract

Plexins receive guidance cues from semaphorin ligands and transmit their signal through the plasma membrane. This family of proteins is unique amongst single-pass transmembrane receptors as their intracellular regions interact directly with several small GTPases, which regulate cytoskeletal dynamics and cell adhesion. Here, we characterize the GTPase Activating Protein (GAP) function of Plexin-B1 and find that a cooperative GAP activity towards the substrate GTPase, Rap1b, is associated with the N-terminal Juxtamembrane region of Plexin-B1. Importantly, we unveil an activation mechanism of Plexin-B1 by identifying a novel functional loop which partially blocks Rap1b entry to the plexin GAP domain. Consistent with the concept of allokairy developed for other systems, Plexin-B activity is increased by an apparent substrate mediated cooperative effect. Simulations and mutagenesis suggest the repositioned JM conformation is stabilized by the new activation switch loop when the active site is occupied, giving rise to faster enzymatic turnover and cooperative behavior. The biological implications, essentially of a threshold behavior for cell migration are discussed.

## Introduction

Plexins are transmembrane receptors with four subfamilies (Plexin-A[1-4], -B[1-3], -C1, -D1) each binding to one or more members of the Semaphorin family of signaling ligands^1^. Semaphorin-Plexin signaling controls cell migration, usually repulsion in neuronal and cardiovascular development^2–3^. Genetic knockout of certain Semaphorins and Plexins can lead to severe neurological development defects, pathogenic angiogenesis, and cancer metastasis^3,4^. Plexin function involves interactions with multiple binding partners such as small GTPases, the co-receptor Neuropilin (which appears obligatory, however only for the Plexin-A family), and diverse receptors or soluble kinases as well as posttranslational modifications and binding to adaptor proteins^3,5^. A unique feature of Plexins, as transmembrane receptors, is the ability to directly interact with Ras- and Rho-family small GTPases via its <u>I</u>ntra-<u>C</u>ellular <u>R</u>egion, ICR, residues 1511-2135^3,6^. The ICR follows the transmembrane region of residues 1487-1510. The ICR consists of an N-terminal intracellular segment, called the Juxta<u>m</u>embrane (JM, 1511-1545) region, <u>G</u>TPase <u>A</u>ctivating <u>P</u>rotein (GAP, 1567-1719, 1916-2135) domain which is split into an N- and C-terminal segments by insertion of a <u>R</u>ho-GTPase <u>B</u>inding <u>D</u>omain (RBD, 1748-1887)^7^ (Fig. 1a). The Plexin GAP domain was shown to stimulate the hydrolysis of GTP to GDP in the Ras GTPase family member, Rap1b^8^. Rap1b, thus deactivated by Plexins, appears to attenuate integrin-mediated cell attachment, which in turn can lead to cell collapse. This is also accomplished via a second function, the binding and activation of a GEF (Guanine nucleotide Exchange Factor) protein for RhoA at a site distinct from the GAP active site and the RBD^9^. The structure of the <u>I</u>ntra-<u>C</u>ellular <u>R</u>egions, ICRs, of different Plexin family members has been solved mostly by x-ray crystallography^10,11^ and unveiled subtly different conformational states of the JM and of a loop which, in the unbound protein protrudes into the Rap1b binding site of the GAP domain (termed “activation segment”).^12^ A dimerization or trimerization of the Plexin ICR (if not of the entire Plexin receptor) is thought to synergize with conformational changes in the JM and activation segment, thereby stimulating its GAP activity^8,12,13^. The ICR of Plexin-A family members can be considerably activated by dimerization in vitro using a coiled-coil or another N-terminal ICR dimerization tag, but likely need additional interactions, perhaps by contacting Neuropilin for full activity. For Plexin-B family members and -D1, while likely present, no enhanced GAP activity was reported for induced dimers, but we note that for Plexin-B1 and –C1, the activity of the wild type ICR is already above that seen in the induced Plexin-A1 dimers^8^. A more dramatic increase of activity is shown in a similarly induced dimerization of zebrafish Plexin-C1^12^, reaching the level of other GAP enzymes. At the outset of this study, we posit that activation may also involve other, not yet characterized segments of the ICR, particularly for the Plexin-B family.

**Figure 1.**
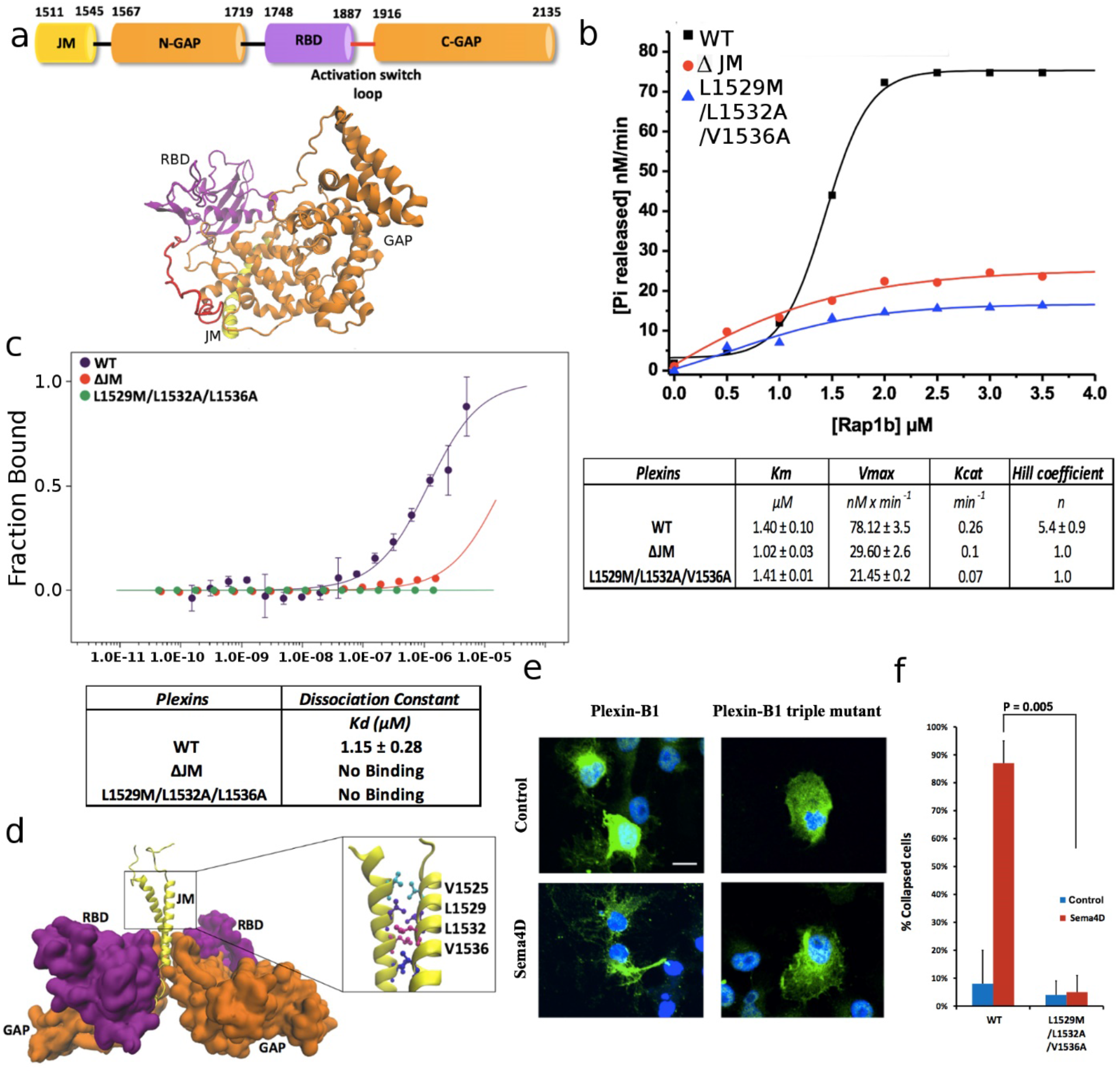
Plexin-B1 activity and evidence of a concentration dependent conformational change via the juxtamembrane region. (a) Domain and main-chain cartoon of the tertiary structure of the Plexin-B1 intracellular region (PDB ID, 3SU8^13^, with missing loops modeled, see M&M). (b) GAP activity of Plexin-B1 (300 nM) with Rap1b as a function of GTPase substrate concentration (0 to 3.5 μM). Enzyme activity and kinetic constants K_M_, V_max_ and k_cat_ were calculated using the Michaelis-Menten equation, cooperativity was calculated using the Hill equation (see M&M). (c) Conformational change assay using Microscale Thermophoresis (MST). The Juxtamembrane domain plays an important role in a Plexin-B1 transition as the triple mutant L1529M/L1532A/V1536A does not show a similar transition. (d) Model of Plexin-B1 dimeric structure, highlighting key JM residues predicted to be part of a coiled-coil^14^. (e) Representative zoom of a slide showing cells in a collapse assay. Cos7 cells were transfected with Plexin-B1 wt (left column) or Plexin-B1 triple mutant L1529M/L1532A/V1536A (right column) and either not treated (top, control) or treated with Sema4D. Scale bar (white) is 10 μm (f) Histogram and significance of a cell count (see Methods). The triple mutation in the JM domain is seen to abolish cell collapse.

## Results & Discussion

### Plexin-B1’s modest GAP function has a cooperative increase in activity at higher substrate concentration

For this project we adopted a fluorescence based *in vitro* assay for the detection of Plexin stimulation of GTP hydrolysis in Rap1b at near physiological protein concentrations, suggested to be around 1 μM or less (Fig. S1) whereas the initial assays of the Zhang laboratory used Plexin concentrations of 10 μM^8^. Using such lower Plexin concentrations, we newly observed a significant activity of Plexin-B1 toward Rap1b, itself depending upon the Rap1b GTPase substrate concentration. The kinetic constants K_M_ (*Michaelis constant*) and V_max_ (maximal rate) were fitted for the Plexin-B1 ICR activity as a function of substrate concentration, in the presence of up to 2 μM Rap1b at 300 nM Plexin-B1 (= E_o_) to be 1.4 ± 0.1 μM and 78.1 ± 3.5 nM/min respectively giving a value for k_cat_ (turnover number = V_max_/E_o_) of 0.26 min^−1^ (Fig. 1b, Fig. S1, M&M Eqn. 1). This turnover number of Plexin-B1 compares with the activity of 0.23 min^−1^ reported for a coiled-coil enforced dimer of Plexin-A1^8^. These GAP activities of Plexin in solution are relatively modest, e.g. Gap1m is 15-fold faster and p120RasGAP even 150 fold faster towards H-Ras (however, the latter with a 100 fold weaker K_M_, compared to Plexin-B1)^14^. The sigmoid or S-shaped behavior of the Plexin-B1 GAP reaction rate as a function of the substrate concentration of Rap1b, with a Hill coefficient of 5.4 ± 0.9 (Fig. 1b, M&M Eqn.2), indicates an enzyme cooperative mechanism within the Plexin-B1 ICR. To our knowledge this is the first time a cooperative mechanism has been described for a GTPase Activating Protein in the literature and the biological implications are discussed below.

### Both the Juxtamembrane segment and a newly discovered loop are responsible for cooperativity

Experiments using Microscale Thermophoresis (MST) as a function of Plexin concentration show a transition with a mid-point at 1.15 ± 0.28 μM for the Plexin-B1 ICR (Fig. 1c, S2). However, in agreement with others^13,15^, even at μM concentration, we failed to observe dimer or trimer formation of the Plexin-B1 ICR by gel filtration and analytical ultracentrifugation, suggesting that a dimer/trimer, if it exists, may be very weak/kinetically labile. At the same time, computer simulations^16^ and a DN-Ara assay in E.Coli (Fig. S3) indicate that the <u>T</u>rans<u>m</u>embrane (TM) helix of Plexin-B1 has a tendency to form a homo-dimer/trimer by itself. The dimerization/trimerization of the TM helices likely further enhances the association of intracellular region for full-length Plexin-B1, considering that the TM helix is immediately followed by a JM helix, which has an intrinsic tendency to form coiled-coiled dimer or trimer^12,13,16^. As a further test, we used cross-linking reagents and could show that Plexin-B1 is more readily cross-linked than the JM triple mutant or ΔJM truncation (Fig. S4). Based on available model structures for TM and JM helices^16^ and given an adequately high sequence identity between human Plexin-B1 and zebrafish Plexin-C1 for those interface residues stabilizing a dimer^17^ (Fig. S5), we built a molecular model of the Plexin-B1 TM+JM dimer with the ICR and examined its stability in 4 independent molecular dynamics simulations of 2 *μ*s each (Fig. S6a,b). In the ICR of Plexin-B1, the dimer would be maintained by hydrophobic contacts established between the side chains of residues F1544, L1547, M1548 from the JM region and L1923, L2036 and L2037 from the GAP domain (Fig. S6c). The N-terminal part of the JM domain is stabilized as a dimer by the putative coiled-coil interactions established between residues V1525, L1529, L1532 and V1536 (Fig. 1d and Fig. S6d). In Fig. S6e, we plot the distribution of orientation angles of the dimeric transmembrane helices as well as dimeric juxtamembrane helices with respect to the membrane normal (see Methods). Both helices are tilted around 20 degrees. Therefore, the dimeric juxtamembrane region has no contacts with the membrane. This directs the GAP and RBD regions away from the membrane and likely better orientates the Plexin-B1 GAP domain for association with the substrate Rap1b.

Based on JM domain mutations of a study of the TM and JM of Plexin-A3^18^, we made mutations of leucine 1529 to methionine, leucine 1532 to alanine, and valine 1536 to alanine, as well as a Juxtamembrane deletion construct, ΔJM. Our GAP activity measurements on both proteins show that removing the JM region of Plexin-B1 abolishes the cooperativity between the Plexin-B1 and Rap1b and also reduced the activity by two thirds (Fig. 1b, table). Similarly, the triple mutant L1529M/L1532A/V1536A, and double mutants L1529M/L1532A or L1532A/V1536A, also lost cooperativity and activity (Hill Coefficient ~ 1, k_cat_=0.07 min^−1^, see also Fig. S6). Interestingly, the deletion/mutations of the Juxtamembrane domain do not affect the Michaelis constant K_M_ of the reaction (Fig. 1b table) suggesting that the GAP active site for Rap1b binding is not affected. No conformational change was detected by MST for both the ΔJM deletion or for the triple mutant protein under our experimental conditions (Fig. 1c).

Treatment with a soluble (N-terminally truncated) ligand, Semaphorin4D (Sema4D) of Plexin-B1 transfected Cos7 cell lines is known to result in the induction of signaling pathways via Plexin-B1: Rap1b: integrin^8,19^. We show in Fig. 1e, that with the triple mutant JM domain, the transfected cells do not collapse compared to the wild type Plexin-B1 after treatment with Sema4D (this is already true for the double mutants, L1529M/L1532A and L1532A/V1536A, Fig. S7). These observations further confirm the role of the Plexin-B1 JM region in a part of its regulatory mechanism. While the JM domain does not significantly affect Rap1b binding, the native JM stimulates the turnover of Rap1b (k_cat_) in the enzyme reaction, especially at higher substrate GTPase concentrations. In order to unveil the underlying molecular mechanism, we examined the structural differences between monomeric and dimeric Plexin-B1 using molecular dynamics simulation. Remarkably, for the first time, we observed a significant structural adjustment of a (so far) unstudied loop in Plexins. For Plexin-B1 this loop is comprised of residues 1887 to 1915 between Plexin-B1 RBD and C-terminal half of the GAP domain. Due to likely intrinsic flexibility, except in one structure of Plexin-B1 bound to Rac1^10^, this loop structure was absent in crystal structures^10,11,12,20^. We noticed that in a monomeric Plexin-B1, the loop extends outward relative to the Plexin-B1 core domain (loop shown in red in Fig. 2a). However, in a model of dimeric Plexin-B1 in our molecular dynamics simulations, the loop moves toward to the JM coiled-coil (Fig. 2b) and is displaced by as much as 15 Å (Fig. S9a). The flexibility of the loop (1887-1915) is compared between the rest state (when the loop is not bound to the JM) and the associated state (the loop is bound to the JM region) (Fig. S9b). Overall, the dynamics of the loop quenches, especially for residues between P1895 and P1905, likely due to the stable association of the loop with the JM. If we were to dock Rap1b to the monomeric apo Plexin-B1 the loop would cause significant steric hindrance to the GAP region, likely affecting Rap1b binding; however, once its conformation is switched to one near the JM coiled-coil region, it provides an open space for Rap1b access. Thus, we expect this loop may play a significant role in modulating the kinetics of the Rap1b enzymatic GTP hydrolysis process. Due to the importance of this loop, as shown below, we name it, the activation switch loop (a.s. loop).

**Figure 2:**
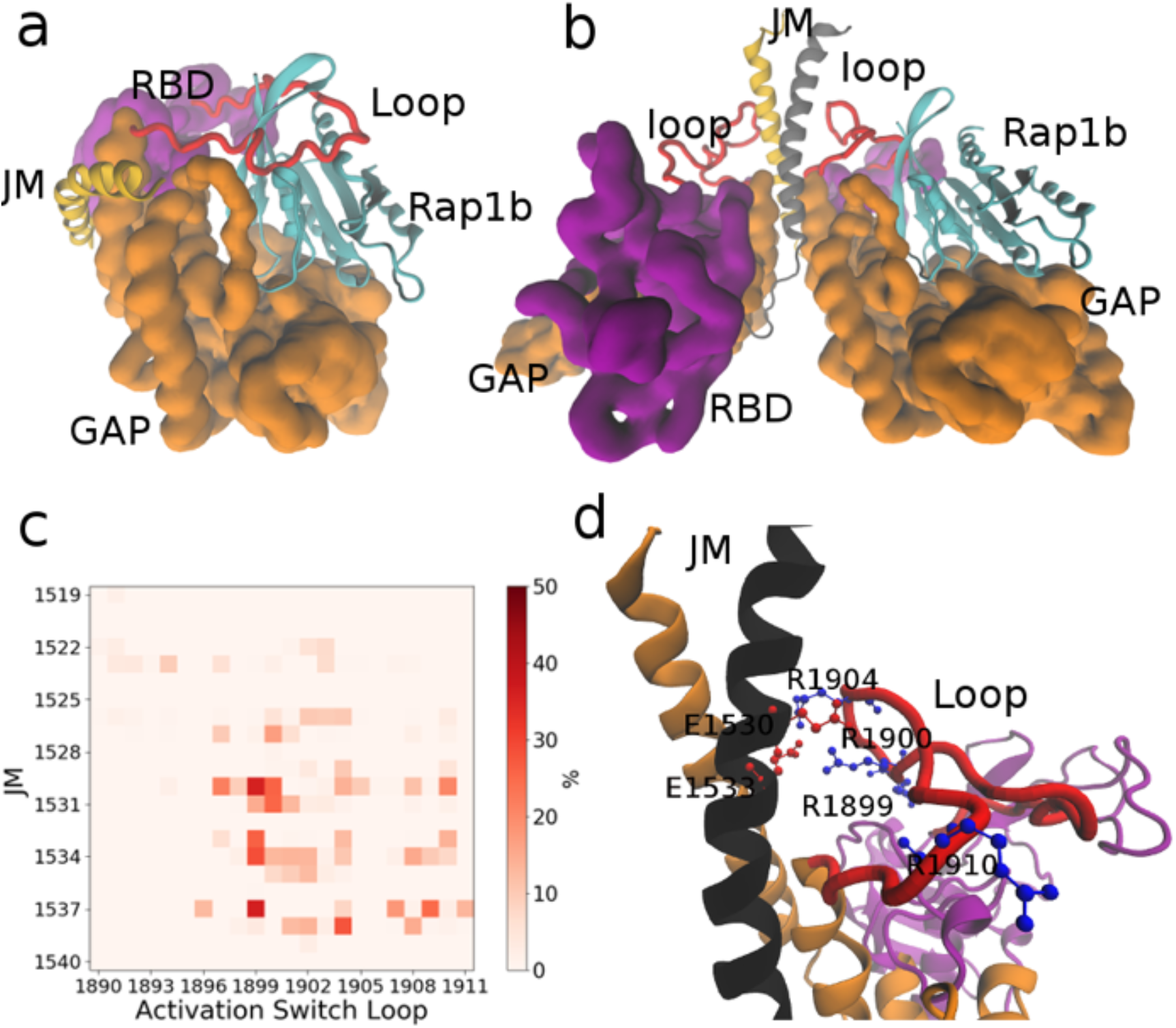
The conformation change of the Plexin-B1 activation switch loop is stimulated by interactions with the JM region. (a) Plexin-B1 monomer (pdb id 3SU8^13^) with loops modeled, as above. The activation switch loop (red) causes a steric hindrance for Rap1b (shown in light blue) entry and/or exit into the GAP domain, here for the monomer. (b) Modeled Plexin-B1 dimer (see M&M). The activation switch loop is displaced and binds to the JM region. (c) Residue-Residue contact maps between the activation switch loop and the JM helix (% of time contact < 4 Å between sidechains as white to red coloring). (d) Highlights of key electrostatic interactions formed between JM helices and the activation switch loop.

Molecular dynamics simulations strongly suggest that the association of the activation switch loop with the JM region is stabilized by the electrostatic interactions between residues R1899, R1900, R1904, R1910 from the activation switch loop and E1530, E1533, D1538 from the JM (Fig. 2c-d), also to a lesser extent by electrostatic pairing with E1907, E1909 and R1537. We mutated positively charged Arginines (res. 1900 and 1904) in the switch loop to negatively charged Glutamic acids, and separately Glutamic acids (res. 1530 and 1533) to Arginines in the JM domain. We then examined the influence of these mutations on the binding between Plexin-B1 and the constitutively active mutant Rap1b.G12V, as well as the GAP activity of Plexin-B1 towards Rap1b (Fig. 3a). Both mutants considerably reduce the binding between Plexin-B1 and Rap1b.G12V (Fig. 3a and Fig. S11). This indicates that the interactions established between the a.s. loop and JM region have a role in limiting the otherwise GAP domain inhibiting conformation of the activation switch loop. In accord with the reduced Rap1b binding affinity, the double mutation E1530R and E1533R in the JM domain, as well as –separately-the double arginine mutation R1900E and R1904E in the activation switch loop, both result in a loss of GAP activity by over 50% (Fig. 3b). An over 50% diminished effect is also seen in the cell collapse assay compared to the wild type Plexin-B1 (Fig. 3c and Fig. S11). We also examined two cancer-associated mutations (given in the COSMIC database, https://cancer.sanger.ac.uk/cosmic): D1538G in the JM and K1523N between the JM and TM region, which, however, have only a small influence on the cell collapse (Fig. S8).

**Figure 3:**
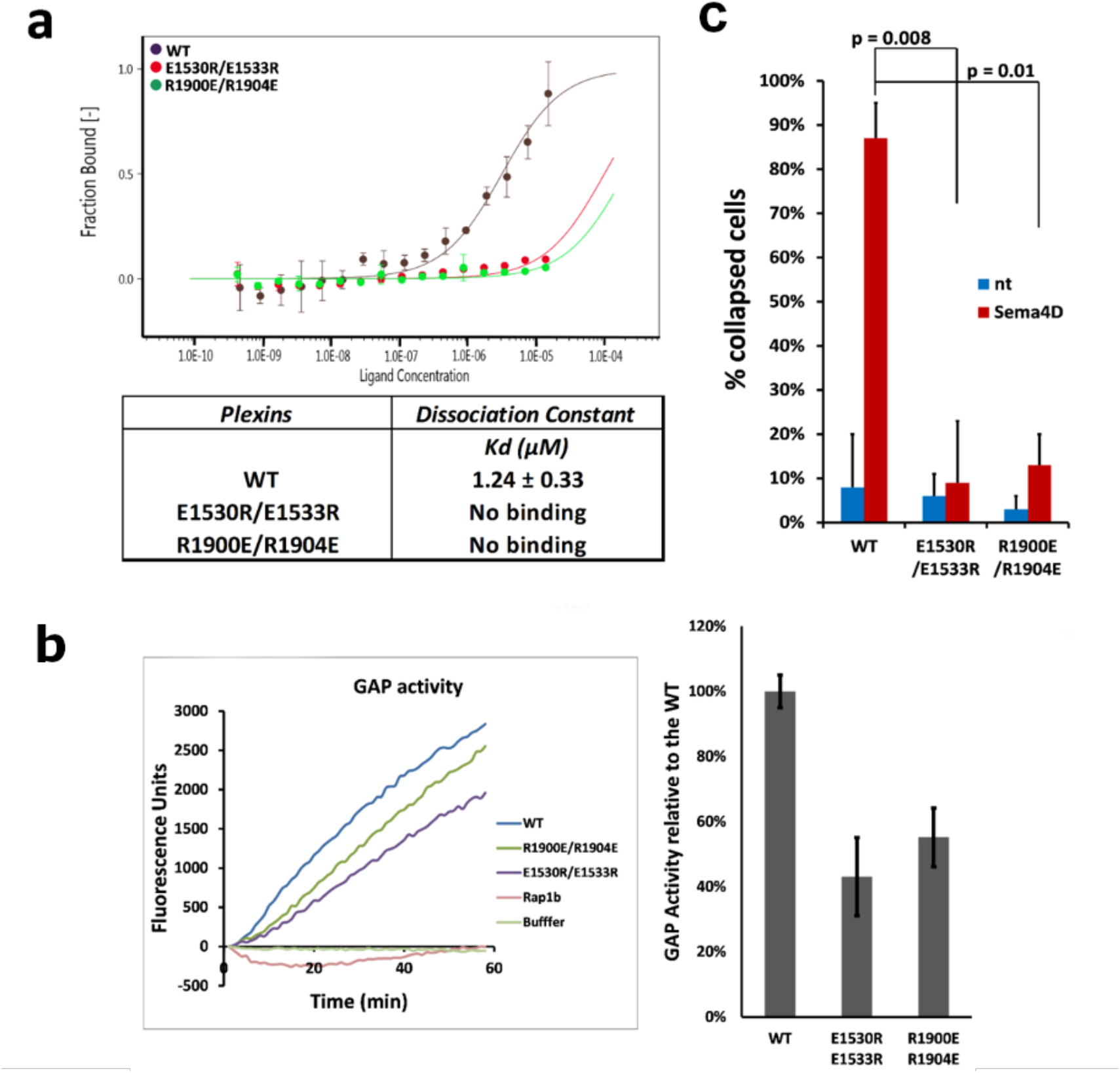
Role of the activation switch loop in Plexin-B1 enzyme activity and in cell collapse function. (a) Plexin-B1: Rap1b.G12V association by MST as a function Rap1b concentration. Glutamic acid 1530 and 1533 in the JM domain and Arginine 1900 and 1904 in the activation switch loop play an important role in the association of Rap1b with Plexin-B1 as no significant binding could be fitted (Kd > 100 μM) when these residues are mutated to Arginine and Glutamic Acid respectively compared to the wild type Plexin-B1. (b) GAP assay and (c) cell collapse assay with Plexin-B1 activation switch loop- and JM mutants.

### Loop-swap chimers show that the function of the activation switch loop is confined to the Plexin-B family

Considering the significance of the activation switch loop, we compared the sequence of this loop between Plexin family members. The loop of Plexin-B1 has an unusual sequence -composed of several Arginines and Prolines- and differs from that of Plexin-B2 and Plexin-B3 (Fig. 4a) but, importantly, the length of the loop is similar within the Plexin-B subfamily. The other Plexin families –A[1-4], -C1, and D1 have shorter loop lengths, with the Plexin-A[1-4] loop being the shortest. We swapped the Plexin-B1 activation switch loop with Plexin-B2 and -B3 loops and with Plexin-A1, -C1 and -D1 loops. For the swap of the Plexin-B1 loop to -B2, -B3, both the Rap1b association and enzyme reaction rate are not, or only slightly, affected (Fig. 4b-c). This is likely because both Plexin-B2 and -B3 have a relatively long loop and have an abundance of basic amino acids similar to Plexin-B1. By contrast, for the switch of Plexin-B1 a.s. loop to the corresponding sequences of Plexin-A1, -C1 and -D1, the enzyme reaction rates are all significantly reduced, while, surprisingly in each case Rap1b binding is stronger (Fig. 4b-c). The results may be explained as on one hand the -A1, -C1 and -D1 loops are shorter and block the active site less (in agreement with stronger binding), but on the other hand the loop is not long enough to reach over to the JM region and be retained there. Such a reaching over may prevent the loop from returning to partially block the GAP region and/or it may further enhance the GAP activity via the previously identified, “activation segment”^11^. There is also the possibility that the switch loop aids the K_off_ rate of product release (weaker Rap1b.GDP binding at the GAP site due to competition from interactions between the activation switch loop and the GTPase).

**Figure 4:**
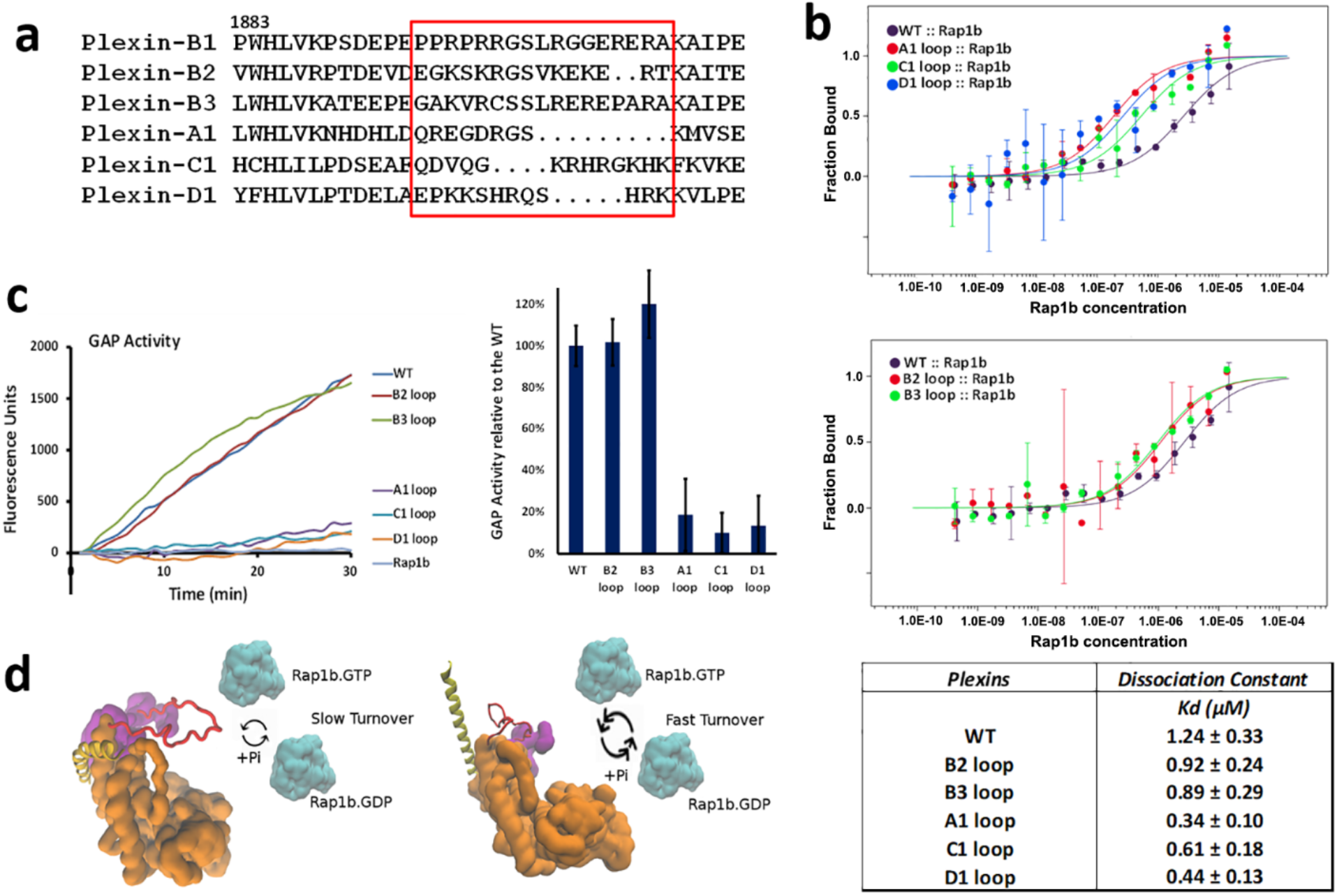
Activation switch loops in different Plexin families. (a) Sequence alignment of activation switch loops (boxed in red) between different human Plexins. Plexin-B1 wild type and activation switch loop swapped constructs, (b) binding with Rap1b.G12V and (c) GAP assay. (d) Model of the allosteric activation process.

### A novel form of kinetic Allostery, Allokairy, arises from a loop movement that competes with substrate

Fig. 4d shows a model of how the activation switch (a.s.) loop alters enzymatic turnover. When near the GAP domain, the a.s. loop acts as a soft obstacle in the process of binding Rap1b.GTP and may also slow the release of Rap1b.GDP. At low Rap1b concentration, the a.s. loop affects binding and release of Rap1b significantly and thus retards the turnover rate or the enzymatic reaction (for example, a low k_cat_ ~0.07 min^−1^ for monomeric Plexin-B1), as the a.s. loop needs to be moved away from the GAP region at every enzymatic turnover process. The steric hindrance of this loop is released upon the formation of interactions with the JM domain, but these are transient, so when the Rap1b.GDP product is released the a.s. loop will return to partially block the active site. At higher substrate concentration the enzyme can achieve a higher reaction rate and thus higher enzymatic activity (k_cat_ = 0.26 min^−1^) because as soon as one product Rap1b is released another substrate will bind, not giving the a.s. loop time to reposition itself near the GAP region – instead it is stabilized away from this region by interactions with the JM. In addition, the cooperativity in the enzymatic reaction could be further supported by a likely slow repositioning of the JM away from and/or towards the GAP region, as suggested by Jones and colleagues^13^, and thus by the virtue of JM - a.s. loop interactions this would also increase the residence time of the activation switch in an open state^13^. The formation of a JM coiled-coil appears to be important in stabilizing the a.s. loop away from the GAP region. This means that at higher Rap1b concentration there is a more frequent turn-over process because both the JM and a.s. loop have not enough time to convert back to states where they partly obstruct the GAP region. Similar inferences of a slow conformational transition which in part are coordinated with an infrequent binding of the substrate at low concentrations and high occupancy at higher substrate concentrations have been made for other systems^21^. The behavior, resulting from a combination of allosteric and hysteretic features has recently been termed “allokairy”.

It should be noted that the proposed model of Fig. 4d can also explain the cooperative behavior within an isolated/homo plexin-B1. However, this scenario is less likely, given that the contacts of the a.s. loop are made with the JM domain of the second protomer. However, dimeric association of intracellular regions is expected to be further enhanced for the full-length receptor at the cell membrane, arising from a concentration effect on limiting diffusion in a large part from 3 to 2 dimensions, as well as from the association of transmembrane (Fig. S3), and extracellular regions. These interactions likely synergize with ICR conformational and activity changes^22^. For example, the dimerization of the Plexin-B1 extracellular region triggered by Sema4D has been proven to be vital the receptors cellular function, since the disruption of the dimeric interface at the extracellular region leads to reduced function of Plexin-B1.^24^ We only have a 4-fold GAP activity increase of Plexin-B1 ICR above the activity of Plexin-B1 ICR ΔJM. This is probably because there are only few associated Plexin-B1 ICRs in solution and even in an associated state the domains may not be orientated optimally to engage the JM, activation- and activation switch loops. For example, the subsequent GAP assay paper with the Zhang laboratory have shown much more dramatic activity increases for Plexin-C1 and –A when the N-terminal coiled-coil – JM domain fusion is made at different spacings to the ICR^12^. With the assistance of the TM, ICR dimer formation would be prevalent. This could result in more stable interactions with the activation switch loop and greatly enhance the activity at the cell membrane.

Our data shows that the extent of activation enhancement due to the activation switch is related to the features of the loop (length as well as charge) of the Plexin-B family of receptors. This finding allows us to speculate on the effect of mutations in the a.s. loop such as R1904W which appears in prostate cancer patients of prostate cancer^23^. In this case, an a.s. loop – JM charge interaction is abolished, also by the placement of a bulky group. Several cancer-associated mutations in the JM region are found in the COSMIC database, such as K1523N, R1537Q, D1538G but do not show an appreciable change in cell collapse in our hands (Fig. S8). We also noticed Serine residue 1902 in the a.s.loop, which is conserved amongst Plexin-B family members. We speculate that this Serine works as a residue site for activation via phosphorylation (the NetPhos3.1 server^24^ predicts phosphorylation by PKA, PKB and PKC kinases). With computational modeling, we studied Plexin-B1 ICR with a phosphorylated Serine 1902 (pS1902) with two simulations each of 0.5 *μ*s. The activation switch loop shows a largely reduced association with the JM coiled-coil (Fig. S12). The pS1902 interacts with the neighboring Arginines 1899, 1904 and 1910, thus greatly reducing the association propensity of the activation switch loop towards JM coiled-coil. The information could be beneficial for the design of possible inhibitors towards Plexin-B1 in cancer treatment, but still yet to be investigated in further detail.

In accord in the plexin receptor being the only example of a transmembrane receptor that harbors a functional GAP domain^1–3^, we may anticipate novel mechanistic features and to our knowledge this the first time plexin has been described as an enzyme whose level of activity experiences cooperativity with respect to the level of substrate concentration. Especially, if this is further amplified near the membrane, it would have profound effects of the function of plexin in regulating cell migration and adhesion as it suggests that at low Rap1b (or possibly other Ras-isoform) GTPase concentrations the active receptor has only a small GAP activity. Once a particular threshold of Rap1b concentration is reaches, here around 1 μM in our in vitro experiments, activity could be dramatically enhanced via the allokairy mechanism, involving the activation switch loop, as described above. This would then lead to a rapid decrease of active Rap1b below threshold and would ensure an all-or none response to the input signal, the cell migration guidance cue, semaphorin. The mechanism would seem especially appropriate for cell collapse (or cell front retraction) as cell attachment via integrins should be switched off completely^2^. While cooperativity in GTPase regulatory enzymes has been described for Guanosine Exchange Factors (GEFs) and is apparent in mechanisms that involve allosteric binding processes to co-activators or membranes^25–27^, intriguingly GTPase Activating Proteins have not been described as cooperative enzymes with respect to substrate concentration so far. To our understanding this possibility has also not been explicitly considered in theoretical models of cell signaling^28,29^, which is already rich in features which modulate the activity of GTPase regulatory proteins and the level of active GTPases.

## Conclusion

In summary, we characterized the GTPase Activating Protein (GAP) function of Plexin-B1 and found that GAP activity towards Rap1b is associated with a cooperative effect, indicating a conformational change in Plexin, such as a repositioning of a newly discovered activation switch loop and of the Juxtamembrane domain. Importantly, this loop of Plexin-B1 plays a profound role in Rap1b turnover (the overall reaction rate of the enzyme) and we confirmed its role by mutagenesis using a GAP activity assay and by cell collapse assays. The effect is specific to Plexin-B sub-family members and thus for the first time suggests subfamily specific differences in the activation mechanism of Plexin intracellular regions.

## Acknowledgements

We thank Drs. Liqun Zhang and Prasanta K. Hota for early work on this project. This work is supported by a NIH R01 grant from the National Eye Institute R01EY029169 and previous grants from NIGMS (R01GM073071 and R01GM092851) to the Buck lab. The Tamagnone lab was supported by the Italian Association for Cancer Research (AIRC-IG grant #19923). Anton Computer time was provided by the Pittsburgh Supercomputing Center (PSC) through Grant R01GM116961 from the National Institutes of Health.

## Author Contribution

Z.L., J.M., and M.B designed the studies, interpreted the data and wrote the manuscript. J.M. conducted cellular experiments, GAP assay and dimerization assays, with L.T. carrying out additional cell collapse assay and contributing to data analysis and manuscript writing revision. Z.L. performed the molecular dynamics simulation. S.K. designed the plasmids, performed the DN-Ara assay and worked on the initial implementation of the GAP assay.

## Methods

### Protein constructs

A series of constructs of the human Plexin-B1 intracellular domain, ICD, were designed and cloned into the vector pCol. Plexin-B1 ICD FL [res. 1511-2135]; Plexin-B1 ICD ΔJM [res. 1565-2135] and site-directed mutagenesis such as L1529M.L1532A; L1532A.V1536A; L1529M.L1532A.V1536A; E1530R.E1533R, R1900E.R1904E; substitution of B1 activation switch loop to the Plexin-B2, -B3, -A1, -C1 or -D1 loop. A series of full-length human Plexin-B1 were also designed and cloned into pCDNA 3.1 for Cos7 cell transfection: Plexin-B1 FL; L1529M.L1532A.V1536A; E1530R.E1533R and R1900E.R1904E. Human Rap1b wild type (gift of Dr. D. Altschuler, University of Pittsburg) [res. 1 to 184] was subcloned into the expression vector pET28c and constitutive active mutant Rap1b G12V was made. All the constructs also contain a His_6_-tag at the N-terminus for affinity purification with Nickel-NTA agarose beads (Qiagen). Site-directed mutagenesis was performed using the QuickChange Lightning kit (Agilent Technology). Soluble Sema4D was obtained from R&D Systems Reagents (see Collapse Assay paragraph below).

### Protein expression and purification

Plexin-B1 constructs were expressed in a soluble form in *E. coli* Rosetta (DE3) (Invitrogen) by induction with 0.2mM Isopropyl-β-D-thiogalactopyraniside (IPTG) at 16°C for 24 hours. Rap1b constructs were also expressed in a soluble form in *E. coli* BL21 (Invitrogen) with 1mM IPTG at 25°C for 16 hours. The cells were harvested at 8,000 *g* for 15 min at 4°C. Cells with expressed Plexin were resuspended in 20 mM Tris-HCl pH 7.5, 500 mM NaCl, 5% (v/v) glycerol, 0.5mM Tris(2-carboxyethyl)phosphine (TCEP), 10 mM Imidazole and 1mg/mL of hen egg white lysozyme. All GTPases were resuspended in 20 mM Tris-HCl pH 7.5, 150 mM NaCl, 4 mM MgCl_2_, 5% (v/v) glycerol, 0.5 mM TCEP. Both buffers were supplemented with 1 tablet per 10 mL of EDTA-free protease inhibitor cocktail (Novagen, Inc.), 1 mM Phenylmethane sulfonyl fluoride (PMSF), 20 μg/mL Leupeptin, 2 μg/mL Antipain, 10 mM/mL Benzamidine, and 50 μg/mL Pepstatin. All resuspended cell pellets were sonicated and then centrifuged at 48,000 g for 1 hour at 4°C. The supernatants were collected and since all the constructs contain a His_6_-Tag at their N-termini, protein purification was accomplished by their affinity to Nickel-NTA agarose beads employing the standard protocol (Novagen, Inc.). There is no evidence that the N-terminal Histidine influences the measurements, thus the his-tag was not cleaved from the proteins. Plexins and GTPases were further purified by either size exclusion chromatography or using a PD-10 Sephadex G-25 desalting column (GE Healthcare) in 20 mM Tris-HCl pH 7.5, 150 mM NaCl, 5% (v/v) glycerol, 2 mM TCEP, 0.005% Tween-20.

### GTP loading

For GAP activity measurement, Rap1b, was loaded with excess GTP right after the purification: 100 μM Rap1b was incubated with 0.5 mM GTP at 30 °C for 10 min in a GTP loading buffer (20 mM Tris-HCl pH7.5, 100 mM NaCl, 5 mM EDTA, 2 mM DTT). The reaction was stopped on ice by adding 50 mM MgCl_2_. To remove unloaded GTP, DTT, imidazole and EDTA from the reaction, the loaded GTPase was purified using a PD-10 Sephadex G-25 desalting column (GE Healthcare) in the final GTPase buffer (20 mM Tris-HCl, 150 mM NaCl, 4mM MgCl_2_, 0.5 mM TCEP, 5% (v/v) glycerol, pH 7.4). For binding experiments, Rap1b was loaded with the very slowly hydrolyzing Gpp(NH)p analogue instead of GTP using the same conditions described above and purified using a PD-10 Sephadex G-25 desalting column in PBS-T 0.1%, (50mM phosphate pH 7.4,150mM NaCl, 0.1% (v/v) Tween-20) supplemented with 4mM MgCl_2_.

### Measurement of GTPase activity in vitro

The Fluorescence-change based GAP assay^30,31^ was performed using a Tecan Infinite M1000 microplate reader (TECAN, Inc.) at 30 °C with proteins in a GAP reaction buffer (20 mM Tris-HCl, pH 7.5, 50 mM NaCl, 0.2 mM TCEP, 2 mM MgCl_2_) and 4 μM of the phosphobinding protein (PBP) (Life Technologies) to detect the inorganic phosphate (Pi) that released from the GTPase following GTP hydrolysis (Fig. S2). The reaction was done in a total volume of 50 μL. First, 0.3 μM of Plexin-B1 was added to the reaction and incubated at 30 °C until the baseline had stabilized (2-3 min). 0 to 4 μM of GTP-loaded Rap1b was then added to the mix and the release of Pi was monitored by the change in fluorescence [using 425 nm (λ_ex_) and 465 nm (λ_em_)] at 1 min intervals for up to 60 min. To calculate the concentration of Pi released in nM/min, a standard curve was measured using 0 to 2.5 μM inorganic phosphate in the same GAP reaction buffer. The enzyme activity and Michaelis constants were calculated from the “steady state” of phosphate release (initial slope where the substrate concentration is in greater excess compared to the enzyme concentration and yield maximum velocities). Kinetic constants K_M_, V_max_ and k_cat_ were calculated using the Michaelis-Menten equation (equation 1) and cooperativity was calculated using the Hill equation (equation 2)

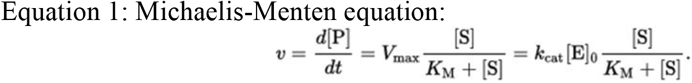

Where

*v* is the reaction rate
[P] is the concentration of product (Rap1b.GDP)formed
*t* is the time of the reaction
[S] is the substrate concentration, Rap1b.GTP
*K*_cat_ is the rate constant
*K*_M_ is the Michaelis constant
*V*_max_ is the maximum rate
[E] is the initial enzyme concentration, the Plexin-B1 ICD or variants at 300 nM.

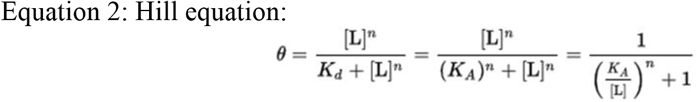

Where:

*θ* is the fraction of the receptor protein concentration that is bound by the ligand
[L] is the free unbound Rap1b concentration
*K_d_* is the apparent dissociation constant derived from the law of mass action
*K_A_* is the Rap1b concentration producing half occupation of the enzyme
*n* is the Hill coefficient

### Measurement of the dissociation constant in vitro

Binding experiments were performed using a Monolith NT.115 Microscale Thermophoresis (MST) instrument (NanoTemper) at room temperature in phosphate buffer saline with 0.1% Tween-20 (PBS-T 0.1%). Proteins are labeled using the NanoTempe Monolith HIS-Tag Labeling Kit RED-trisNTA which labels His-tags with a fluorescent group. 40 nM of each labeled Plexin construct was mixed with a serial dilution of unlabeled GTPases or Plexin-B1 in 0.2 mL micro reaction tubes (NanoTemper) and then transferred to premium capillaries (NanoTemper). Microscale thermophoresis measures the change of the diffusion of molecules in microscopic temperature gradients upon protein ligand binding/protein-protein association. The dissociation constant K_d_ was obtained by fitting the binding curve with the quadratic solution for the fraction of fluorescent molecules that formed the complex between proteins A and T, calculated from the law of mass action K_D_ = [A]*[T]/[AT] where [A] is the concentration of free fluorescent molecule and [T] the concentration of free titrant and [AT] the concentration of complex of A and T.

### DN-Ara assay

Plasmid pAraGFPCDF was first transformed in the AraC deficient *E. coli* strain SB1676. Plasmids for the DN-Ara^32^ assay were a gift from Dr. B.W. Berger. The cells containing this plasmid were then made competent on media containing 50μg/mL Spectinomycin and co-transformed with the plasmids pAraTMwt and pAraTMDN. The cells containing the three plasmids were selected on LB plates containing 50μg/mL Spectinomycin, 50μg/mL Ampicillin and 25μg/mL Kanamycin. One colony from each plate was grown in 5mL selective LB media at 37 °C for 18 hours at 250 rpm. 50μL of culture from each sample was then transferred to 5mL selective LB media and grown again for 18 hours using the same conditions. Cell cultures were then diluted in the same media to 0.6 OD_600nm_ and induced with 1mM IPTG for 6 hours at 37 °C and 250 rpm. A 2-fold dilution was made in LB and 100μL of culture form each sample was then transferred to a black 96-well, clear bottom (Greiner) plate. Measurements at 560nm for cell density and GFP fluorescence at 585nm (excitation) and 530nm (emission) were collected using a M1000 Tecan plate fluorescence reader. The relative fluorescence was calculated as the ratio of the fluorescence at 530nm/fluorescence at 560nm. The results were then divided by the negative control pGFPPDF/pTrc99/pTrc-RSF empty vectors.

### Cell collapse assay.^33^

1μg of recombinant Plexin-B1 FL WT pCDNA 3.1 plasmid or Plexin-B1 FL mutants pCDNA 3.1 (as described above) was transfected to 80% confluent COS 7 cells on coverslips using 3 μL Trans IT 2020 transfection reagent (Mirus Bio) for 48 hours at 37 °C. The cells were then treated with 25 nM soluble Sema4D (R&D Systems Reagents) for 35min or not treated (control). The cells were then fixed and treated with PB1 C-terminal anti Rabbit as primary antibody and Goat anti-rabbit IgG H&L (Alexa Fluor 488, green) as secondary antibody. The nuclei were stained with DAPI (blue). After mounting the coverslips on glass, the results were acquired by digital images using a microscope equipped with a confocal imaging system. The counts are the morphometric quantification of collapsed cells. As done by others^33^, we have only counted the positively receptor-expressing cells, and for each experiments we examined multiple slides to assess at least 50 cells expressing the plexin. The collapsed cells are defined as rounded cells having a diameter ≤ 15 μm.

### Molecular dynamics simulation

Molecular Models of the Plexin-B1 was constructed using the available Plexin-B1 structure (PDB entry 3hm6 and 3su8). Missing loops were built with Modeller.^34^ Dimeric Plexin-B1 was modeled by sequence alignment and structure superimposition with dimeric Plexin-C1 of zebrafish (PDB entry 4m8n). The Plexin-B1 transmembrane helix was embedded into the membrane of 680 POPC and 170 POPS with the CHARMM-GUI.^35^ The protein-membrane system was solvated by TIP3P water with 150 mM NaCl for a total of ~450,000 atoms. Four independent simulations were performed for dimeric Plexin-B1.

All simulations were performed with NAMD/2.12^36^ package for initial equilibration of 50 ns. The simulations were then transferred to the Anton 2 supercomputer for production of 2.0 μs. The CHAMRM36m^37^ force field was used in the simulations for water and biomolecules. The van der Waals (vdW) potential was cut off at 12 Å and smoothly shifted to zero between 10 and 12 Å. The electrostatic interactions were calculated with the Particle-Mesh Ewald (PME) method with a real-space cut-off of 1.2 nm. The SHAKE algorithm was applied for all covalent bonds to hydrogen. A time step of 2 fs was employed and neighbor lists updated every 10 steps. The temperature was coupled by to a Langevin thermostat of 310 K, whereas the pressure control was achieved by a semi-isotropic Langevin scheme at 1 bar. After an equilibration time of 50 ns the trajectories were transferred to Anton2^38^ for calculation of the μs long trajectories. In order to calculate the orientation of a helix dimer relative to the membrane, the alpha carbons of the first four residues of the helix A and helix B are selected to calculate the center of mass of the N-terminus. Similarly, the last four residues of each two helices are used to calculate the average position of the C-terminus. The directional vector of a helix dimer is then the line connecting the N-terminal center to the C-terminal center. The orientation angle of the transmembrane or juxtamembrane helix dimer is then calculated by measuring the angle between the membrane normal direction and the helix dimer directional vector. The trajectories were analyzed with VMD^39^ and with scripts for standard analysis.

