## supporting_information for "Plexin-Bs enhance their GAP activity with a novel activation switch loop generating a cooperative enzyme"

[1]Department of Physiology and Biophysics, Case Western Reserve University, School of Medicine, 10900 Euclid Avenue, Cleveland, Ohio 44106, U. S. A. [2]Faculty of Physical, Biological and Mathematical Sciences, Catholic University, Rome, Italy. [3]Department of Pharmacology; [4]Department of Neurosciences, and [5]Case Comprehensive Cancer Center, Case Western Reserve University, School of Medicine, 10900 Euclid Avenue, Cleveland, Ohio 44106, U. S. A.

+ equal contributions

**Figure S1: GAP activity raw data of plexin-B1 ICR WT (a), and mutants ΔJM (b), L1529M/L1532A/V1536A (c), in presence of increasing amounts of GTP-loaded Rap1b.** Upon GTP hydrolysis to GDP, the release of inorganic phosphate Pi – captured by the Phosphate Binding Protein (PBP) – is monitored over time by the change of fluorescence at 425nm (λex) and 465nm (λem). Enzyme activity was calculated from the initial slope (steady state) of phosphate release where the substrate concentration is in greater excess compared to the enzyme concentration and yield maximum velocities.

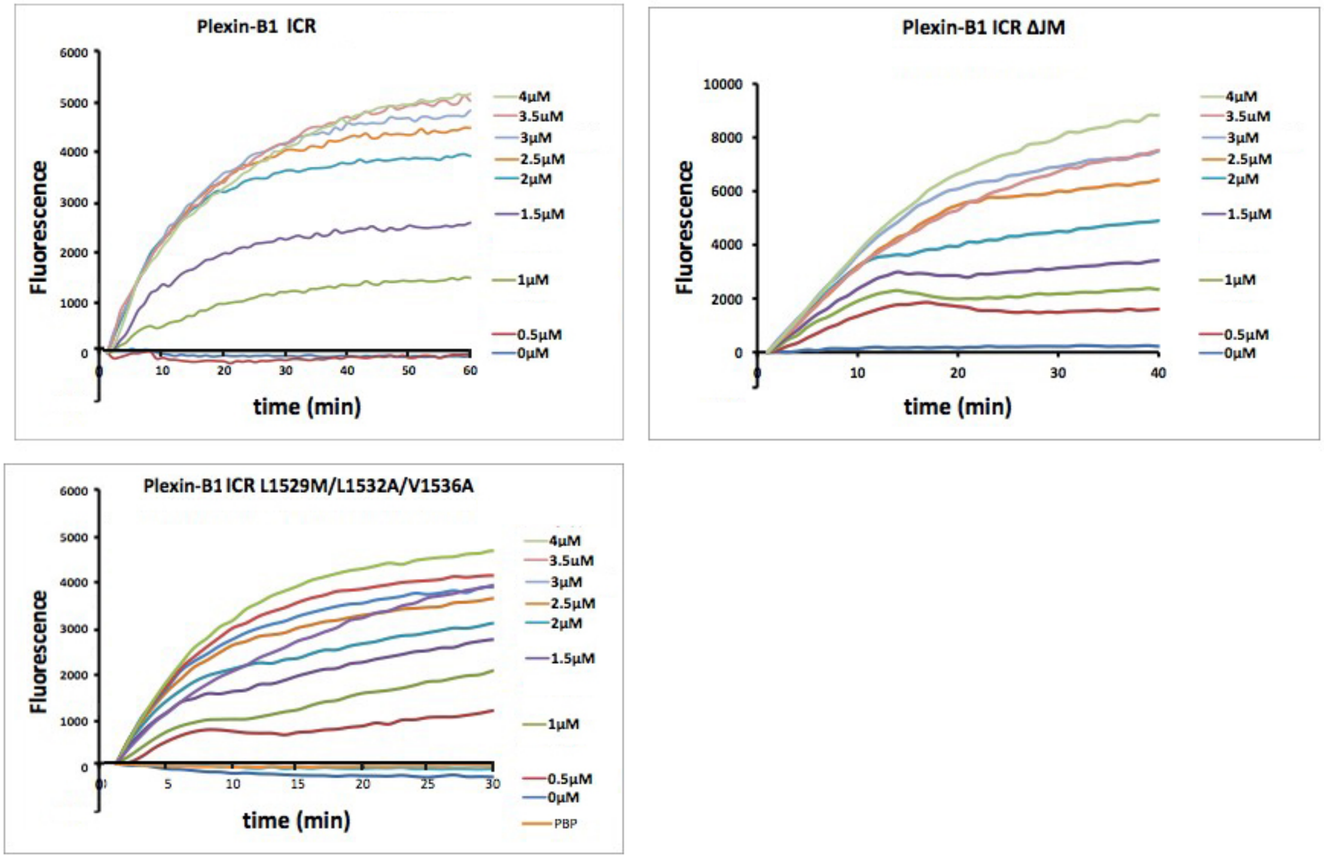

**Figure S2: Representative Microscale Thermophoreis (MST) traces** upon adding 40 nM His-tag fluorescently labeled plexin-B1 WT (a), ΔJM (b), and L1529M/L1532A/V1536A (c), with increasing concentration of the same proteins unlabeled. The microscale thermophoresis measures the change of the diffusion of molecules in microscopic temperature gradients upon protein/protein association.

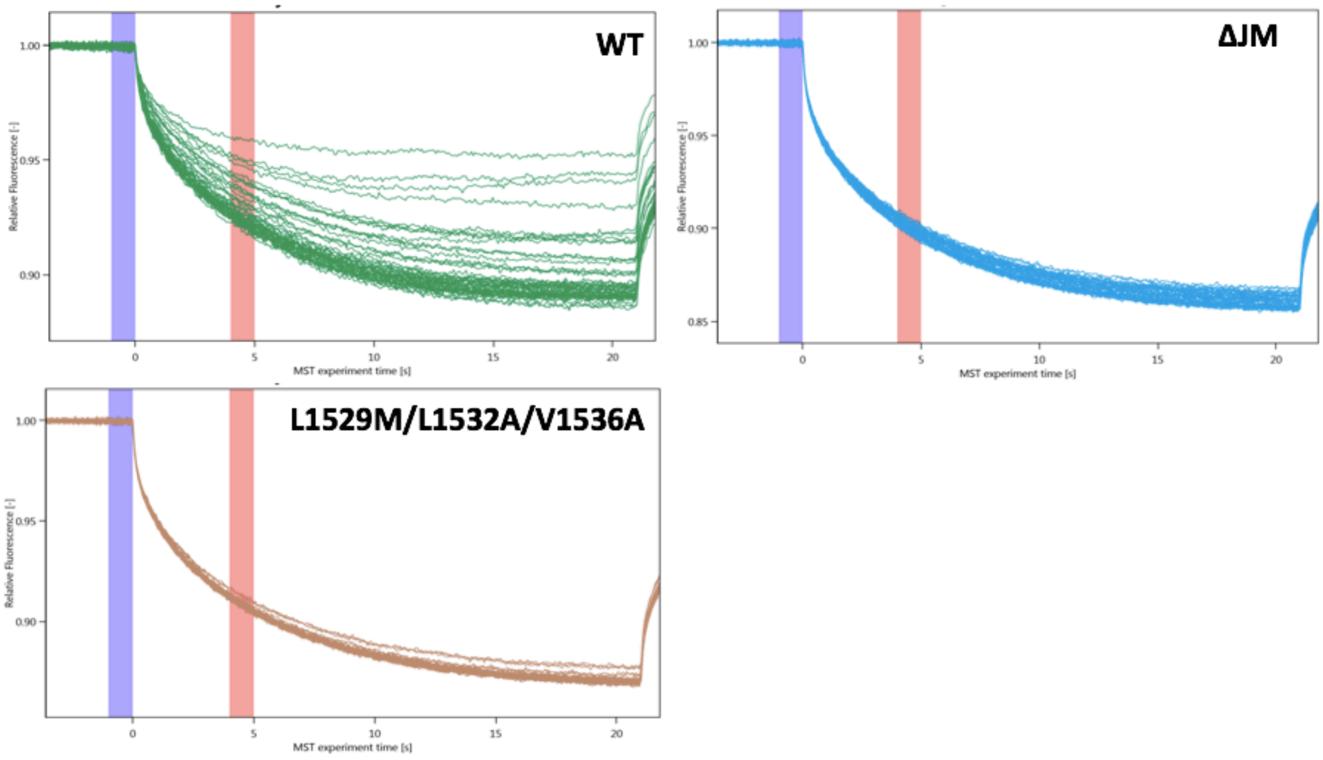

**Figure S3: DN-Ara assay of transmembrane dimer of Plexin-B1 transmembrane helix.** Plexin-B1 transmembrane helix was inserted into the pAra TMwt vector labled as Plexin-B1 and tested for its homodimerization in AraC deficient *E. coli* using the DN-Ara assay. The normalized fluorescence of 6.2 was compared to the positive control experiment using Integrin with a normalized fluorescence of 7.8. Plexin-B1 TM helix was also inserted into pAra TMDN vector (arabinose dominant negative) labeled as Plexin-B1*and was compared to pAra TMDN vector with Integrin (Integrin*) experiment. Both experiments gave similar results of 0.9 and 0.8 respectively. Plexin-B1pAra TMwt (Plexin-B1) and Plexin-B1pAra TMDN (Plexin-B1*) were also both inserted in AraC deficient *E. coli* and tested. The normalized fluorescence of 6.2 was similar to Integrin/Integrin* with a value of 4.1. All the experiments were normalized to the GFPpCDF/pTrc99/pTrc-RSF empty vectors.

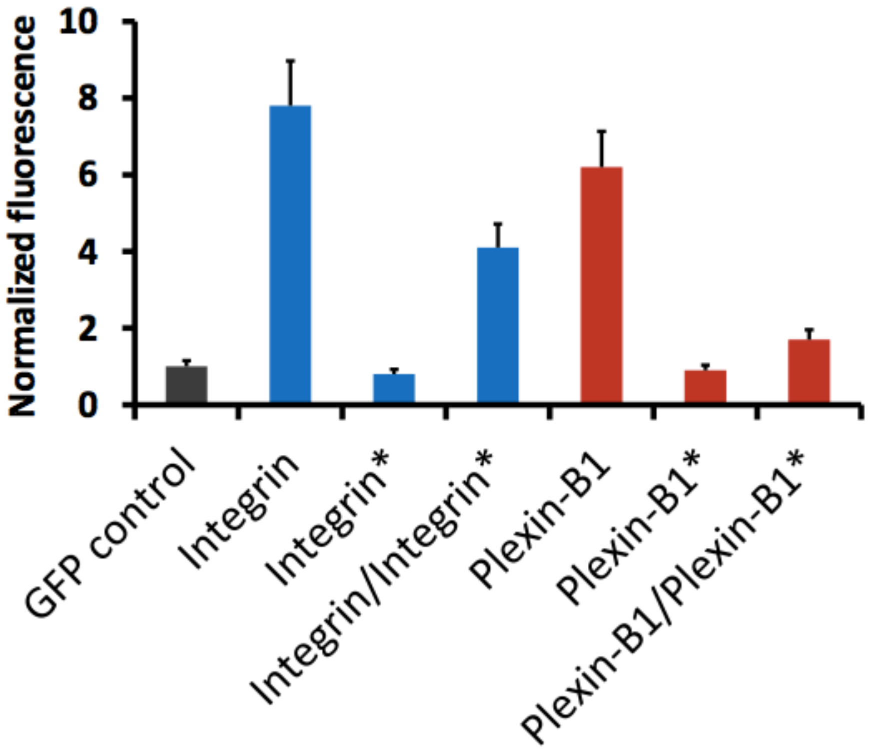


| | Normalized Fluorescence |
|---|---|
| GFP control | 1.0 |
| Integrin | 7.8 |
| Integrin* | 0.8 |
| Integrin/Integrin* | 4.1 |
| Plexin-B1 | 6.2 |
| Plexin-B1* | 0.9 |
| Plexin-B1/Plexin-B1* | 1.7 |

**Figure S4: Chemical cross-linking of PB1-RBD, PB1wt and PB1 ΔJM.** Cross-linking of a control protein – the plexin-B1 Rho GTPase Binding Domain, (RBD) - using 10µg and 50µg of protein and 0, 0.05, 0.1 and 1mM DDS. Incubation was 10 min at 37 ºC. After the incubation, the samples were directly heated to 100ºC with Loading dye and run onto a 16% SDS-PAGE gel. Figure S4a. shows the results of the cross-linking using these conditions. For the experiment with Plexin-B1wt ICR, 10 and 50µg of the protein were incubated using the same conditions as above and the protein was run on a 10% SDS-PAGE after the cross-linking reaction. Then 50 µg of Plexin-B1wt and ΔJM were cross-linked using the same conditions as above. a) RBD after cross-linking reaction at two different concentrations. It is clear that at both protein concentrations, a considerable amount of dimer is seen and that at 1 mM DDS cross-linking reagent. b) Plexin-B1WT ICR at 10 and 50µg of the protein concentration with various levels of cross-linking reagent DDS. c) Cross-linking data for plexin-B1 WT ICR (left) and –B1 WT ICR ΔJM (right). Both proteins at 16 µg (however, ΔJM seems to show more intense staining). Still, the FL ICR appears to dimerize/multimerize earlier than the ICR ΔJM protein.

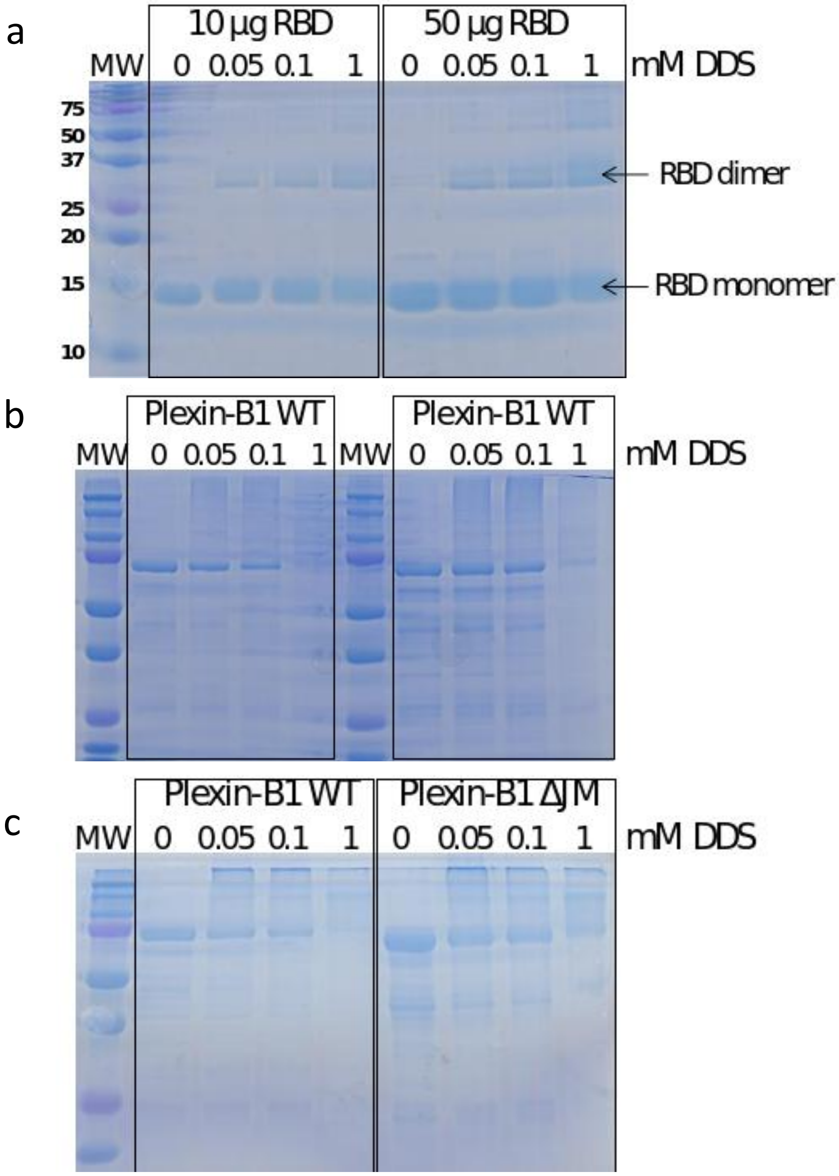

**Figure S5: Sequence alignment between Plexin-B1 (human), Plexin-C1 (zebra fish) and Plexin-A3.** The sequence identity between the JM regions (residues 1520-1550 [1-31 below] for Plexin-B1,) is 65% for plexin-B1 compared to –A3 and 67% for plexin-B1 compared to plexin-C1.

```
Plexin-C1      -MSKKMNDQLELMESNIRRDIRQGFVDLQTEKSDLIDNV--GAIPFLDYKHFASRIFFPE    57
Plexin-B1      RDYKKVQIQLENLESSVRDRCKKEFTDLMTEMTDLTSDLLGSGIPFLDYKVYAERIFFPG    60
Plexin-A3      RTLKRLQLQMDNLESRVALECKEAFAELQTDINELTNHMDEVQIPFLDYRTYAVRVLFPG    60
                  *::: *:: :** :    :: *.:* *: .:* ..:    ******: :* *::**

Plexin-C1      AGTLTAVMIRDIGED-SEQTTVDEKCLAFAELIRDKQFLSCFVHALEEQKNFSIKDKCTV    116
Plexin-B1      HRES--PLHRDLGVPESRRPTVEQGLGQLSNLLNSKLFLTKFIHTLESQRTFSARDRAYV    118
Plexin-A3      IEAH--PVLKELDTP----PNVEKALRLFGQLLHSRAFVLTFIHTLEAQSSFSMRDRGTV    114
                      : :::.       .*::    :.:*:..: *:  *:*:** * .** :*:  *

Plexin-C1      ASLLTLALHGDLLYLTEIMEDLLQSLMDQSS--NANPKLLLRRTESIVEKLLTNWMSICL    174
Plexin-B1      ASLLTVALHGKLEYFTDILRTLLSDLVAQYV--AKNPKLMLRRTETVVEKLLTNWMSICL    176
Plexin-A3      ASLTMVALQSRLDYATGLLKQLLADLIEKNLESKNHPKLLLRRTESVAEKMLTNWFTFLL    174
               ***  :**:. * * * ::. ** .*: :      :***:*****::.**:****::: *

Plexin-C1      YGFLRESVGQPLFLLVSALTQQISKGPVDSVTEKALYTLSEDWLLCQAQDFEPLKLKVVF    234
Plexin-B1      YTFVRDSVGEPLYMLFRGIKHQVDKGPVDSVTGKAKYTLNDNRLLREDVEYRPLTLNALL    236
Plexin-A3      HKFLKECAGEPLFLLYCAIKQQMEKGPIDAITGEARYSLSEDKLIRQQIDYKTLTLHCVC    234
               : *:::..*:**::*   .:.:*:.***:*::* :* *:*.:: *: :  ::. *.*: :

Plexin-C1      AVGTGEEISESLEVIALTCDTIQQVKEKILQTFQRKFGFRYTQQIRDIEIEYEKE-GKFV    293
Plexin-B1      AVGPGAGEAQGVPVKVLDCDTISQAKEKMLDQLYKGVPLTQRPDPRTLDVEWRSGVAGHL    296
Plexin-A3      ---PENEGSAQVPVKVLNCDSITQAKDKLLDTVYKGIPYSQRPKAEDMDLEWRQGRMTRI    291
                       :  : * .* **:* *.*:*:*: . : .       . . :::*:..     :

Plexin-C1      MLQEVDDTSEIRGHVTMLNTLKHYQVGDGACIKVITPKIHAPLKT--------------    338
Plexin-B1      ILSDEDVTSEVQGLWRRLNTLQHYKVPDGATVALVPCLTKHVLREN-------------    342
Plexin-A3      ILQDEDVTTKIECDWKRLNSLAHYQVTDGSLVALVPKQVSAYNMANSFTFTRSLSRYESL    351
               :*.: * *:::.     **:* **:* **: : ::

Plexin-C1      -------------QNSVKDDKNFSIKYFHLVDPDIDTDLS--------NHPEKKALKIKE    377
Plexin-B1      -----QDYVPGERTPMLEDVDEGGIRPWHLVKPSDEPEPPRPRRGSLRGGERERAKAIPE    397
Plexin-A3      LRTASSPDSLRSRAPMITPDQETGTKLWHLVKNHDHADHREGDR---------GSKMVSE    402
                                :   .: . : :***.   . :                :  : *

Plexin-C1      MYLIKLLSTKVAVHSFVENLFKSIWGLPN--NKAPLAVKYFFDFLDEQAERKKITDPDVL    435
Plexin-B1      IYLTRLLSMKGTLQKFVDDLFQVILSTS---RPVPLAVKYFFDLLDEQAQQHGISDQDTI    454
Plexin-A3      IYLTRLLATKGTLQKFVDDLFETVFSTAHRGSALPLAIKYMFDFLDEQADQRQISDPDVR    462
               :** :**: * ::::.**::**: : .         ***:**:**:*****::: *:* *.

Plexin-C1      HIWKTNSLPLRFWVNILKNPDFVFSDMEKSPHLDGCLSVIAQAFMDSFSLTDTHLDKHSP    495
Plexin-B1      HIWKTNSLPLRFWINIIKNPQFVFD-VQTSDNMDAVLLVIAQTFMDACTLADHKLGRDSP    513
Plexin-A3      HTWKSNCLPLRFWVNVIKNPQFVFD-IHKNSITDACLSVVAQTFMDSCSTSEHRLGKDSP    521
               * **:*.******:*::***:***. :...   *. * *:**:***: : :: :*.:.**

Plexin-C1      TNKLLYGKDIPQYKQEVKSYYKLVKDQTSISSQELKTFLQEESKKHQNEFNESAALRELY    555
Plexin-B1      INKLLYARDIPRYKRMVERYYADIRQTVPASDQEMNSVLAELSWNYSGDLGARVALHELY    573
Plexin-A3      SNKLLYAKDIPNYKSWVERYYRDIAKMASISDQDMDAYLVEQSRLHASDFSVLSALNELY    581
                *****.:***.**  *: **  : . .  *.*::.: * * *  : .::.   **.***

Plexin-C1      KYMQRYFTEIFQKLEQTDAPSNLKE--NMHRVKELFDNMKRSGWN    598
Plexin-B1      KYINKYYDQIITALEEDGTAQKMQLGYRLQQIAAAVENKVTDL--    616
Plexin-A3      FYVTKYRQEILTALDRDASCRKHKLRQKLEQIISLVSSDS-----    621
                *: :*  :*:  *:.  :  : :  .:.::  ...
```

**Figure S6: Computational Modeling of Plexin-B1 JM-ICR dimer at a POPC/POPS membrane.** (a) Initial structure shown as ribbon representation. (b) Initial configuration of Plexin-B1 dimer modeled at the membrane. Color scheme for Plexin-B1 is the same as Fig. 1a. POPC in grey and POPS in green. (c) RMSD of the Plexin-B1 ICR dimer at the membrane for four independent simulations. Residue-Residue contact maps between (d) JM (protein A) and GAP domain (protein B), and (d) JM (protein A) and JM (protein B). (e) Distribution of orientation angle of dimeric tramsmembrane helix and dimeric juxtamembrane helix. The last 1500 ns of the four independent simulations are used for the analysis.

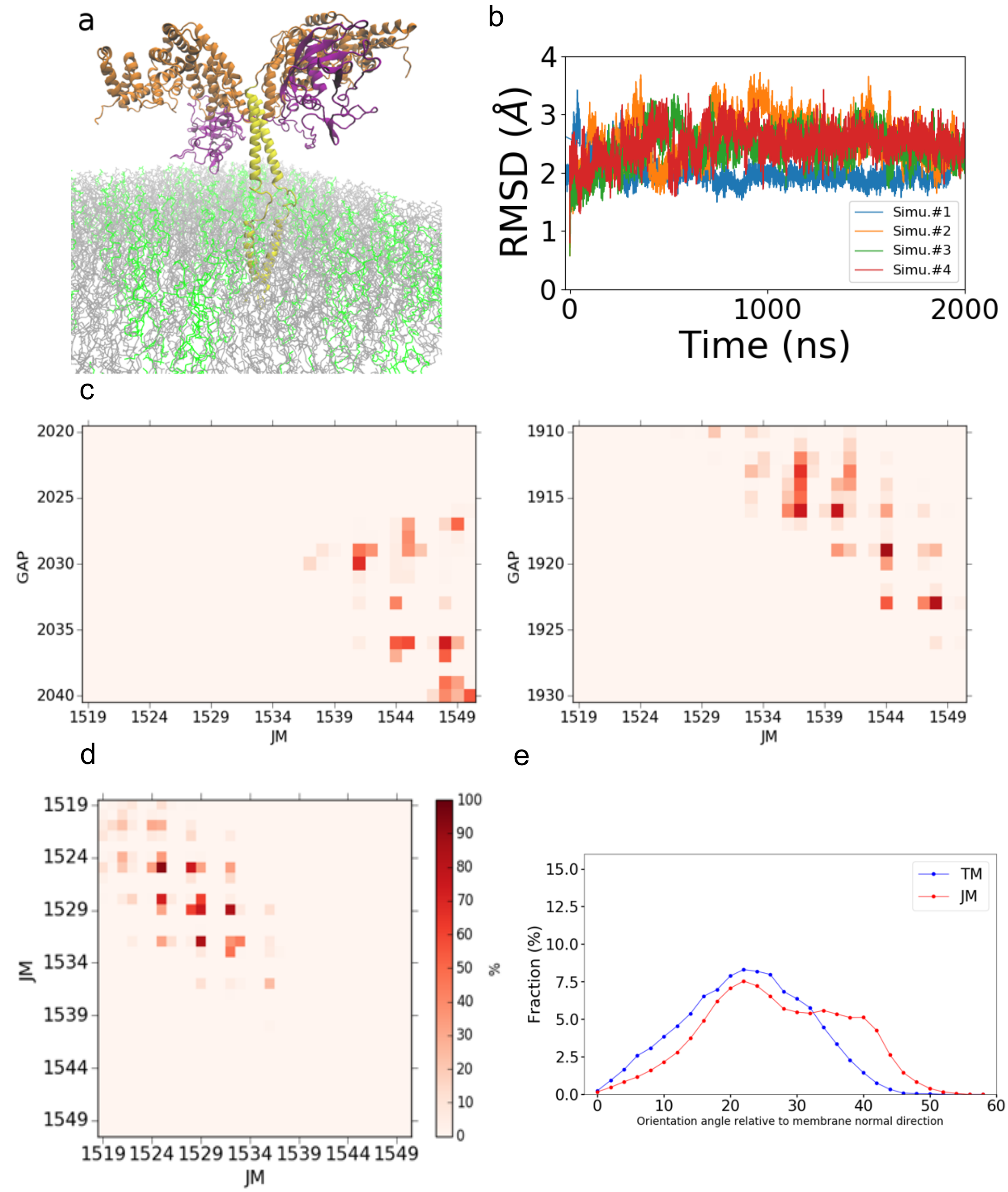

**Figure S7: GAP activity measurements.** Fluorescence traces for plexin-B1 ICR L1529M/L1532A (a), and L1532A/L1536A (b) in presence of increasing amounts of GTP-loaded Rap1b. (c) Initial reaction rate plotted as a function of substrate concentration.

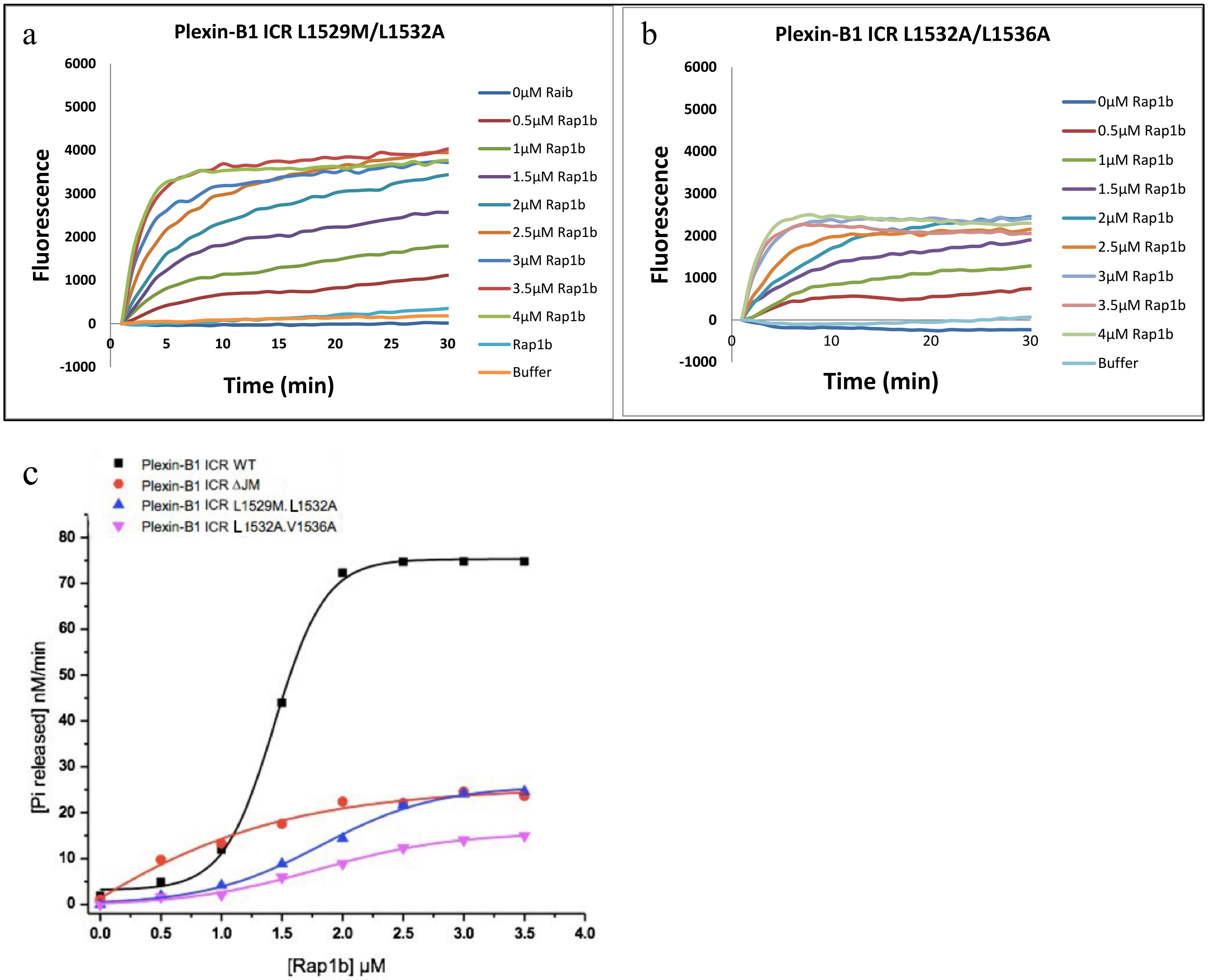

**Figure S8: Cell collapse assay.** 1 μg of Plexin-B1 FL WT or mutants were transfected to COS 7 cells and either treated with 25 nM soluble Sema4D for 35 min or not treated (control). The cells were then fixed and treated with PB1 C-terminal anti Rabbit as primary antibody and Goat anti-rabbit IgG H&L (Alexa Fluor 488) as secondary antibody. The nuclei were stained with DAPI. After mounting the coverslips on glass, the results were acquired by digital images using a microscope equipped with a confocal imaging system. Collapsed or non-collapsed cells were determined based on the reduced surface area compared to the control cells.

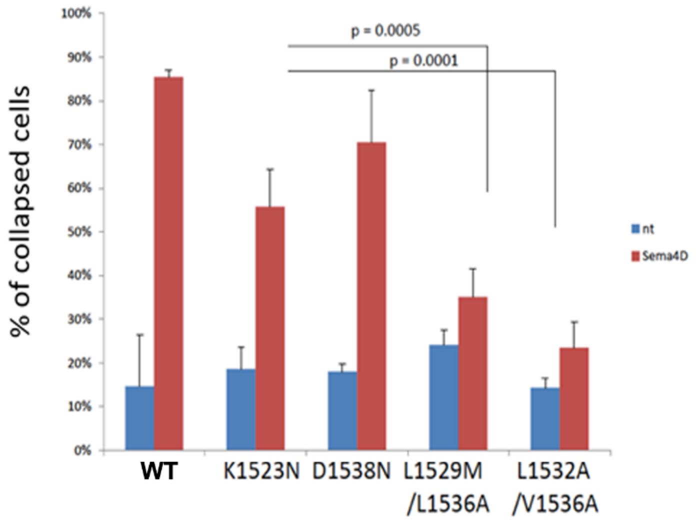

**Figure S9: Computational study of activation loop.** (a) Displacement of activation switch loop of protein A to the JM region of protein B in four independent simulations. The residues 1899-1910 are chosen for the activation switch loop; and the residues 1530-1533 are chosen for the JM helix. (b) RMSF of the activation switch loop (1887 to 1915) in the rest state (not bound to the coiled-coil) and in the associated state (displaced and bound to the coiled-coil). The last 1500 ns of the four independent simulations are used for the analysis.

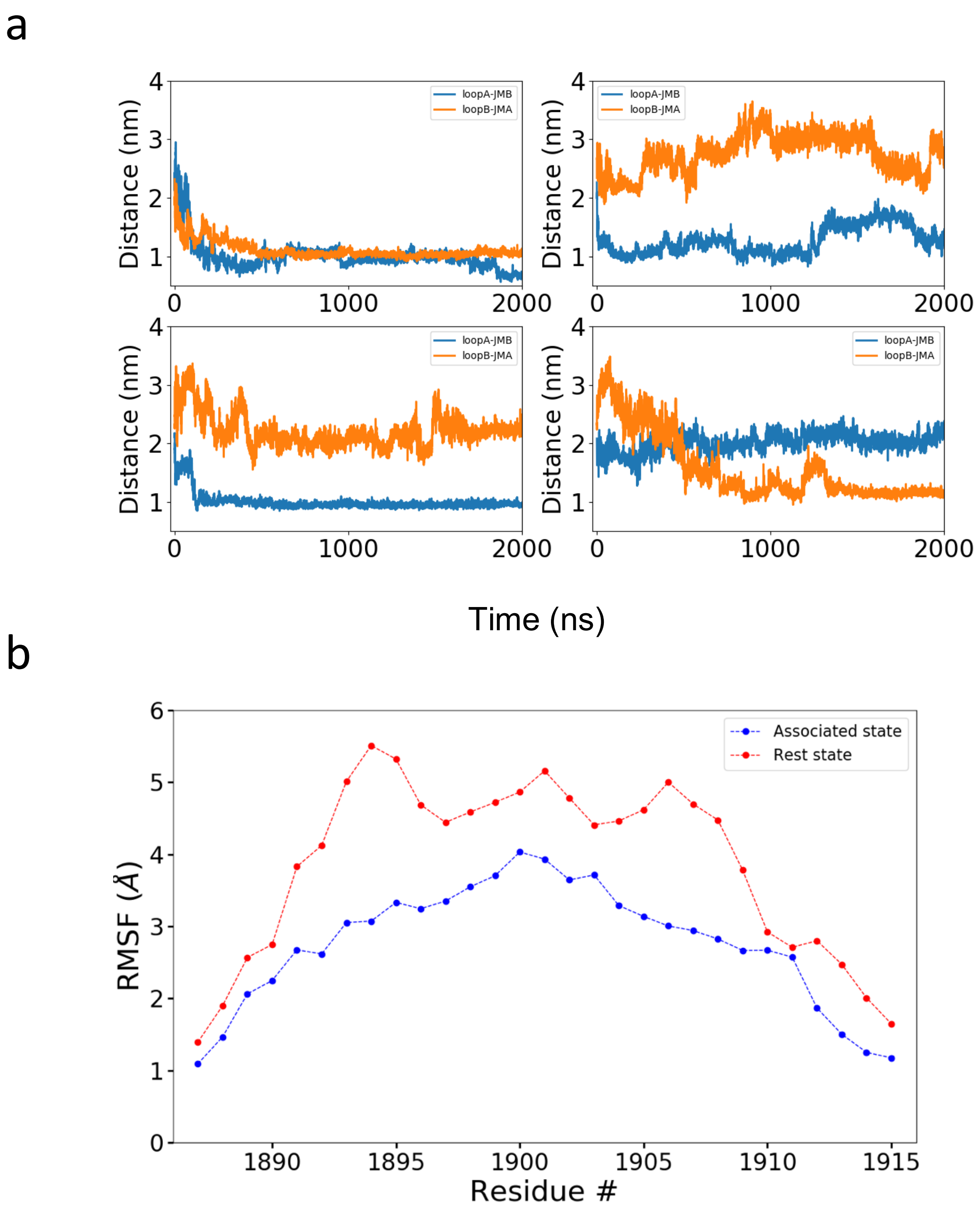

**Figure S10: MST of Plexin-B1 conformational change**, apparent $K_D$, for loop R1900E/R1904E and JM E1530R/E1533R mutations).

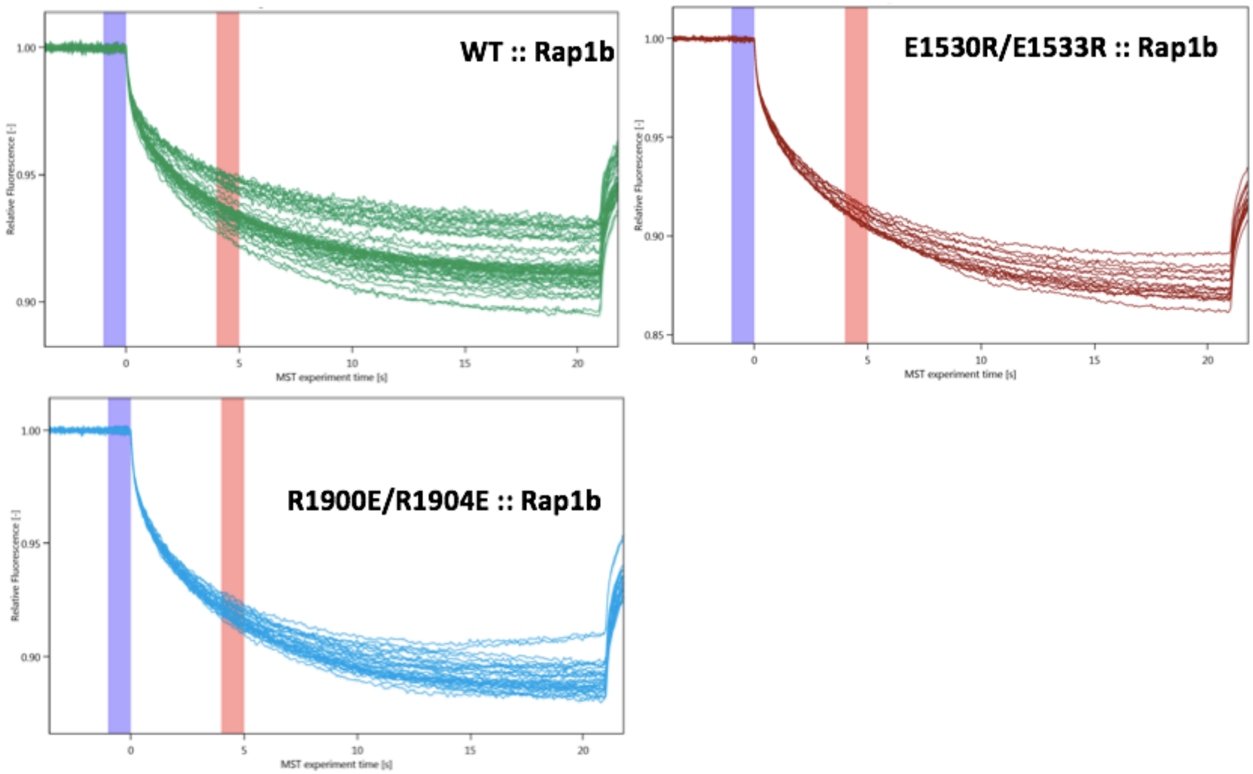

**Figure S11: Cell collapse assay for JM/loop mutations.**

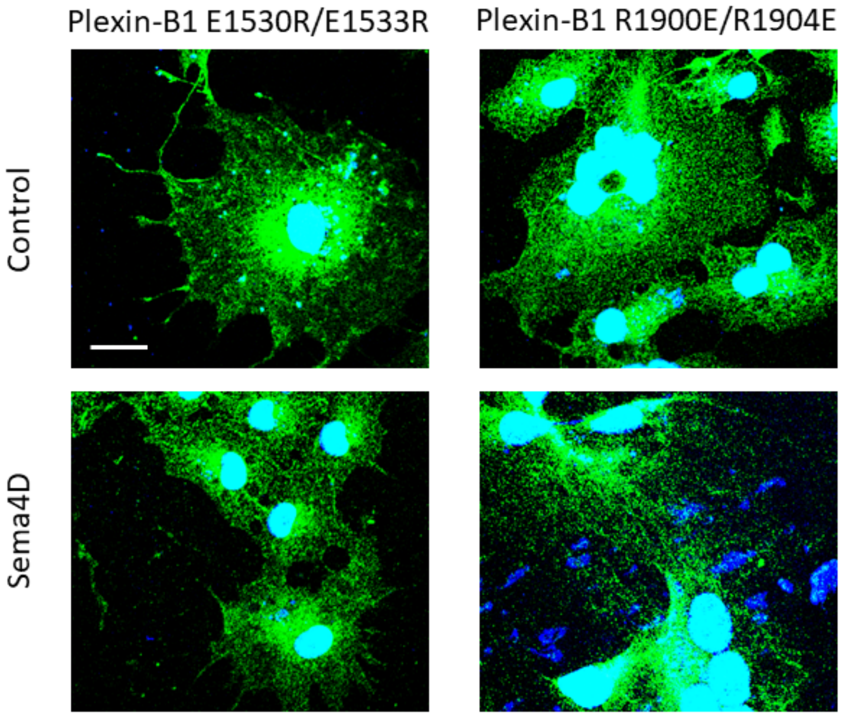

**Figure S12: Computational modeling of Plexin ICR with pS1902.** (a) Displacement of activation switch loop of protein A to the JM region of protein B in two independent simulations. (b) Activation switch loop with phosphorylated serine 1902. Highlight interactions of the pS1902 with neighboring Arginines.

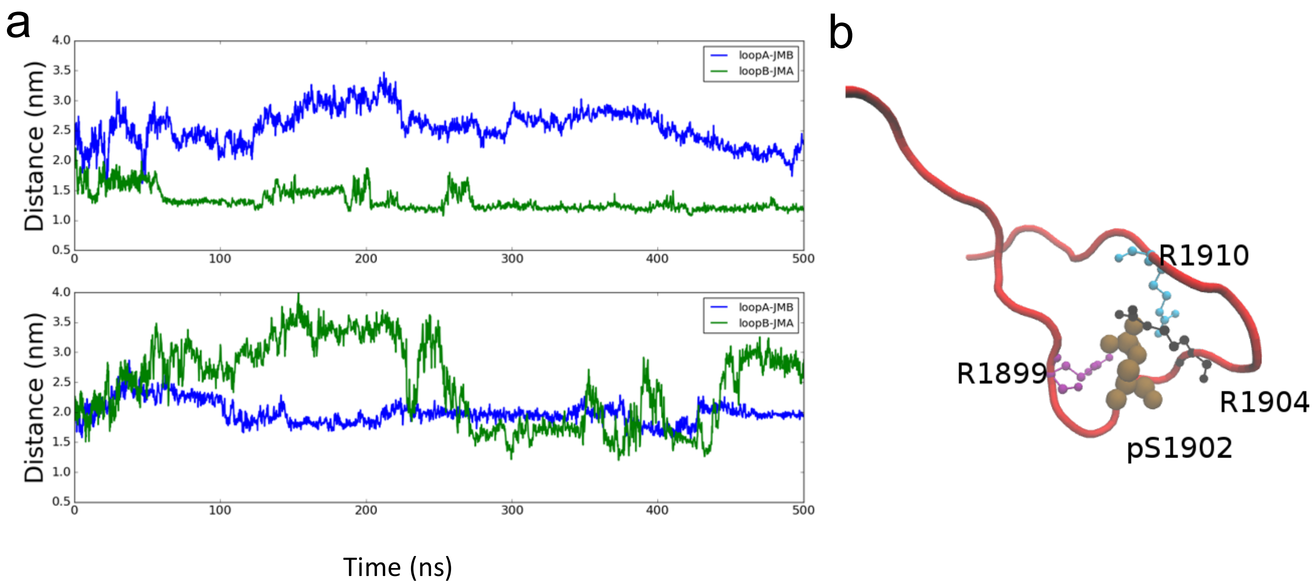
